# High-precision photoacoustic neural modulation uses a non-thermal mechanism

**DOI:** 10.1101/2024.02.14.580331

**Authors:** Guo Chen, Feiyuan Yu, Linli Shi, Carolyn Marar, Zhiyi Du, Danchen Jia, Ji-Xin Cheng, Chen Yang

## Abstract

Neuromodulation is a powerful tool for fundamental studies in neuroscience and potential treatments of neurological disorders. Both photoacoustic (PA) and photothermal (PT) effects have been harnessed for non-genetic high-precision neural stimulation. Using a fiber-based device excitable by a nanosecond pulsed laser and a continuous wave laser for PA and PT stimulation, respectively, we systematically investigated PA and PT neuromodulation at single neuron level. Our results show that the laser energy needed for PA neurostimulaion is 1/40 of that needed for PT stimulation to achieve the same level of cell response recorded by Ca^2+^ imaging. The threshold energy for PA stimulation is found to be further reduced in neurons overexpressing mechano-sensitive channels, indicating the direct involvement of mechano-sensitive channels in PA stimulation. Electrophysiology study of single neurons upon PA and PT stimulation was performed by patch clamp recordings. Electrophysiological features stimulated by PA are distinct from those induced by PT, confirming that PA and PT stimulations operate through distinct mechanisms. These insights offer a foundation for rational design of more efficient and safer non-genetic neural modulation approaches.

## Introduction

Neuromodulation with high spatial precision is a valuable tool for understanding the flow of information in the nervous system and treatment of neurological disorders. Non-electromagnetic neuromodulation developed in the past two decades has added many options to this toolkit. Insights gained on the cellular mechanism of these non-electrical methods deepen our understanding of neuroscience and facilitate rational design of methods for clinical applications. For example, optogenetics allows control over the activity of selected cells using a combination of genetic engineering and light. Some promising clinical potential for optogenetics is emerging, for example, in treating retinal degenerative disease ^1^. However, broader clinical use is limited, as it requires genetic manipulation ^2^. Focus ultrasound has been demonstrated as a non-genetic non-invasive brain modulation method and has been tested in multiple clinical trials. Emerging evidence suggests that mechanosensitive ion channels on the neuron membrane are involved in ultrasound stimulation ^3^. Yet its spatial resolution is typically a few millimeters.

Non-genetic optical stimulation offers a promising solution to address abovementioned limitations. Infrared neurostimulation uses the photothermal (PT) effect to trigger neuronal activity ^4^. Strong light-absorbing nanomaterials, including gold nanoparticles ^5^, Si nanowires ^6^, polymers ^7^ and carbons ^8^, were used to enhance the thermal effect. Two main PT stimulation mechanisms have been suggested. The first is the direct thermal mechanism, where a temperature increase of a few degrees, induced by light, could activate thermosensitive ion channels ^9^ [shelley]. Yet, the substantial increase in temperature raises significant safety concerns, particularly in clinical applications. The second is the opto-capacitive mechanism, which requires a rapid and transient temperature increase at speed of kilokelvins per second upon light irradiation. This modulates the capacitance of the cell membrane ^10, 11^, and drives a sufficient transmembrane capacitive current ^12,^. Yet, such steep slope of temperature increase precludes its application in vivo due to the challenges to focus a pulsed laser through scattering tissues.

Recently, the photoacoustic (PA) effect has been harnessed as a new approach for high-precision neuromodulation ^13^. In 2020, Jiang et al. reported the development of a fiber optoacoustic emitter ^14^ capable of sub-millimeter neurostimulation both in vitro and in vivo. In addition to the fiber device, other modalities were also developed for different applications, including biocompatible optoacoustic films for neural regeneration ^15^, photoacoustic nanotransducers ^16^ for single cell stimulation, and optically generated focused ultrasound for transcranial brain stimulation ^17^.

Despite these technical advances, the mechanism of PA stimulation remains intriguing. Physically, in a PA process, absorption of pulsed light results in a local and transient heat, inducing a thermal expansion of the absorber and leading to a propagating mechanical wave at the ultrasound frequencies ^18^. When the PA emitters are placed nearby the targeted neurons, the neurons are expected to simultaneously sense the pressure wave from the PA effect and the transient temperature rise due to the heat generated by the PA emitters. Notably, the energy conversion efficiency of a typical photoacoustic process is less than 3% ^18, 19^. Therefore, it is crucial to dissect which processes play a major role in triggering the neural activity and to find out the mechanism that boosts the efficiency in PA neuromodulation.

In this study, we first demonstrate a universal fiber-based device that can be used for both PT and PA stimulation at single-neuron level. The device acts as a PT or PA emitter driven by corresponding laser conditions. On the recording side, while conducting simultaneous whole-cell patch clamp recordings is highly challenging in neuronal stimulation by transducer ultrasound, it is feasible upon fiber-based PA and PT stimulation. With this tool, we systematically investigated and compared PT and PA neuromodulation under different laser conditions. The results reveal that the laser energy required for achieving the same level of activation in PA stimulation is approximately 1/40 of that needed for PT stimulation. Studies on temperature changes in neurons show that PA stimulation is associated with negligible temperature rise, confirming its non-thermal mechanism. Ruling out the thermal effect, molecular mechanisms for PA neuromodulation were further studied. Through overexpressing and pharmacologically blocking of ion channels, mechanosensitive ion channels including TPRC1, TRPP2 and TRPM4 were identified to play significant roles in boosting the stimulation effect.

## Results

### A fiber device for PA and PT stimulation

In order to directly compare the PA and PT stimulation at single cell level, we designed a fiber emitter (FE) that allows both conditions by coupling it to different excitation lasers. Specifically, the tip of a commercial optical fiber with a diameter of 200 μm was coated with a layer of candle soot, as an absorber, and a second layer of polydimethylsiloxane (PDMS), as described previously ^20^. According to the PA theory ^21^, under irradiation by nanosecond pulsed laser, both thermal and stress confinement conditions are met for efficient PA conversion. The same FE was connected to a nanosecond (ns) pulsed laser (RPMC One 100uJ-1030nm) for PA stimulation and to a continuous wave (CW) laser (Cobolt Rumba, 1064 nm) for PT stimulation (**Fig. 1a**).

**Figure 1.**
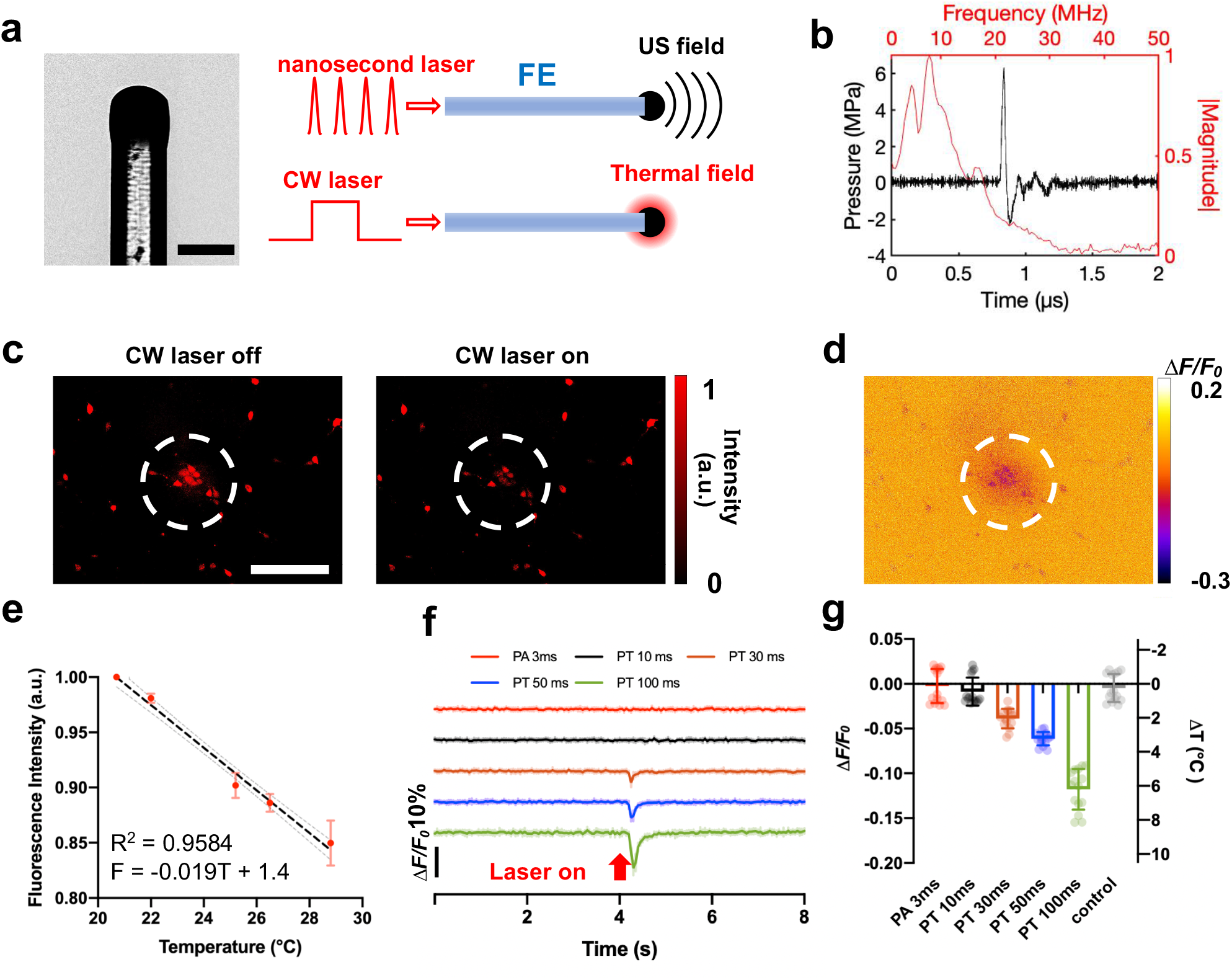
FE used for both PT and PA stimulation and its thermal effect measured in the neurons labelled with mCherry. **a**. Picture and schematic of FE generating photoacoustic and photothermal signal using different lasers. Scale bar: 200 μm. **b**. Representative photoacoustic signal generated by FE using the nanosecond pulsed laser. Black: wave form in the temporal region. Red: frequency spectrum. Laser energy: 45 μJ per pulse. **c**. Representative mCherry fluorescence imaging of neurons under FE before and after CW laser is on. Laser condition: 1064 nm CW laser, 120 mW, 100 ms duration. Scale bar: 200 μm. **d**. Contrast image showing the decrease of fluorescence intensity of the same view field in c. Dashed circles: area of the FE. **e**. Calibration curve of the normalized mCherry fluorescence intensity change due to temperature increase. Standard deviation (SD) was taken from 5 neurons. **f**. Average mCherry fluorescence traces taken from 15 neurons under the FE applied with different laser conditions. Laser condition for PA: the nanosecond laser, 120 mW, repetition rate 4.2 kHz. Laser condition for PT: the CW laser, 120 mW. **g**. Average minimum *ΔF/F*_*0*_ of the 15 mCherry labelled neurons in f and estimated temperature change under different conditions. Control: no laser.

Under the pulsed laser condition, the FE efficiently emits a PA signal (**Fig. 1a)** measured by a needle hydrophone with a diameter of 40 μm (Precision Acoustic, UK). The hydrophone was aligned with the FE and placed at 50 µm away from the tip of the FE (**Fig. S1**). Under the laser condition of 45 µJ per pulse, the FE produced an ultrasound waveform with a peak-to-peak pressure of over 8 MPa (**Fig. 1b**), which is consistent with previously published results and is sufficient for efficient PA neurostimulation ^20^. When coupled with the CW laser, the FE functions as a pure PT source and generates a very localized thermal field (**Fig. 1a**) visualized by a thermal camera (**Video S1**). Together, owing to the strong absorption given by the candle soot and the high expansion coefficient of PDMS ^22^, the FE serves as a point source of ultrasound and heat under specific laser conditions.

### Neuronal temperature under PT and PA conditions

To directly measure the thermal effect of the FE on neurons under PA and PT conditions, we monitored the temperature change on neurons using the fluorescence of mCherry reported as a sensitive temperature indicator ^3^. mCherry was delivered to neurons via AAV9 viral vector transfection (Addgene, pAAV-hSyn-mCherry). Through fluorescence imaging, we confirmed the reduction in fluorescence from mCherry when temperature increases. As depicted in **Fig. 1c-d**, when applying the CW laser with a duration of 100 ms and an average power of 120 mW to the FE, the fluorescence intensity of mCherry exhibited a decrease of over 10%.

To quantify the relation between the fluorescence decrease and the actual temperature increase, we performed a calibration experiment by imaging mCherry-labeled neurons at different temperatures. The temperature of the neurons was raised from 20°C to 30°C in a controlled manner using a dish heater and monitored by a thermal coupler in the medium. For each temperature, the medium was heated until the temperature stabilized at the targeted temperature over 20 s. **Fig. 1e** plots the normalized fluorescence intensity of mCherry as a function of temperature measured by the thermal coupler. For each data point, five cells in the field of view were selected and the fluorescence intensity of the cell was determined by averaging all pixels from the cell. The fluorescence excitation light was only turned on after the temperature stabilized and was turned off immediately after capturing the fluorescence image to minimize any possible photobleaching that might affect the fluorescence intensity. A linear fit was applied to calibrate the intensity of the mCherry fluorescence to the temperature. The results indicate that the fluorescence intensity decreased by 1.9% for every 1°C increase in temperature (**Fig. 1e**), consistent with previously published results ^3^.

Based on the calibration, we evaluated the temperature change on the mCherry-labeled neurons (N=15) under a wide range of laser conditions used for PA and PT stimulation. The FE was placed at a controlled distance of less than 50 μm away from a neuron. Temperature changes were calculated based on the fluorescence change of mCherry and the calibration curve. Under the FE with a 10 ms duration pulse of the CW laser at a power of 120 mW, no significant fluorescence change was observed, and thus the temperature increase is negligible (**Fig. 1f** black line, -0.01 ± 0.02°C). For PT conditions with longer laser durations, a significant drop in fluorescence intensity was observed upon laser activation. Specifically, with the CW laser durations of 30, 50, and 100 ms, the corresponding temperature increase was calculated to be 2.0 ± 0.6°C, 3.2 ± 0.4°C and 6.2 ± 1.2°C (**Fig. 1g** orange, blue, and green), respectively.

We also performed mCherry temperature measurements under PA conditions. Based on previously reported studies ^20^, the PA signal generated by a burst of nanosecond pulsed laser of 3 ms is sufficient to activate neurons. Here, maintaining the average laser power at 120 mW, the same as the PT laser condition, a pulse train (1030 nm, 3 ns, RPMC, Fallon, MO, USA) of 3 ms duration was applied to the FE and the fluorescence intensity of mCherry was monitored (**Fig. 1f**, red line). Minimal temperature increase of 0.00 ± 0.02°C was observed (**Fig. 1h**), comparable to the control conducted with no laser. Notably, the fluorescence imaging of mCherry was conducted with a 20 Hz imaging rate. While this imaging speed was insufficient to precisely track the transient heating associating with the ns laser pulses or the fast rising of the temperature during the 3 ms laser duration, the cooling process was sufficiently slow to be recorded through the fluorescence imaging. Notably, the slow decay is indeed presented in the PT groups shown in **Fig. 1g**. Collectively, by the temperature measurements under different laser conditions, the PA groups showed a clear difference from the PT groups. Specifically, the temperature rise on the neurons is negligible under the pulsed laser condition.

### PA stimulation uses much lower laser energy than PT stimulation

To determine the energy dosage needed for PA and PT stimulation, we deployed calcium imaging to monitor the neural activities under different laser conditions. Although widely used to monitor neuronal activity, GCaMP was reported to show a high sensitivity to temperature, which consistently resulted in significant thermal artifacts during PT experiments ^23^. Thus, we opted to employ OGD-488 (Oregon Green 488) as a calcium indicator. OGD-488 exhibits lower sensitivity to thermal influences and allows more accurate observation of neuronal activity.

Firstly, we applied the 1030 nm pulsed laser with an average power of 120 mW to an FE placed within 50 μm away from the neuron. When the laser duration was 1.5 ms, corresponding to five 3-ns laser pulses, a transient stimulation was observed (Max Δ*F*/*F*_0_ =11.7 ± 4.3%) (**Fig. 2a**, pink trace). When the laser duration was increased to 3 ms, i.e. ten laser pulses, the responses from the neurons were stronger and the averaged fluorescence change reached 13.9 ± 2.3%. Fluorescence images can be found in **Supplementary Figure S2a, b**.

**Figure 2.**
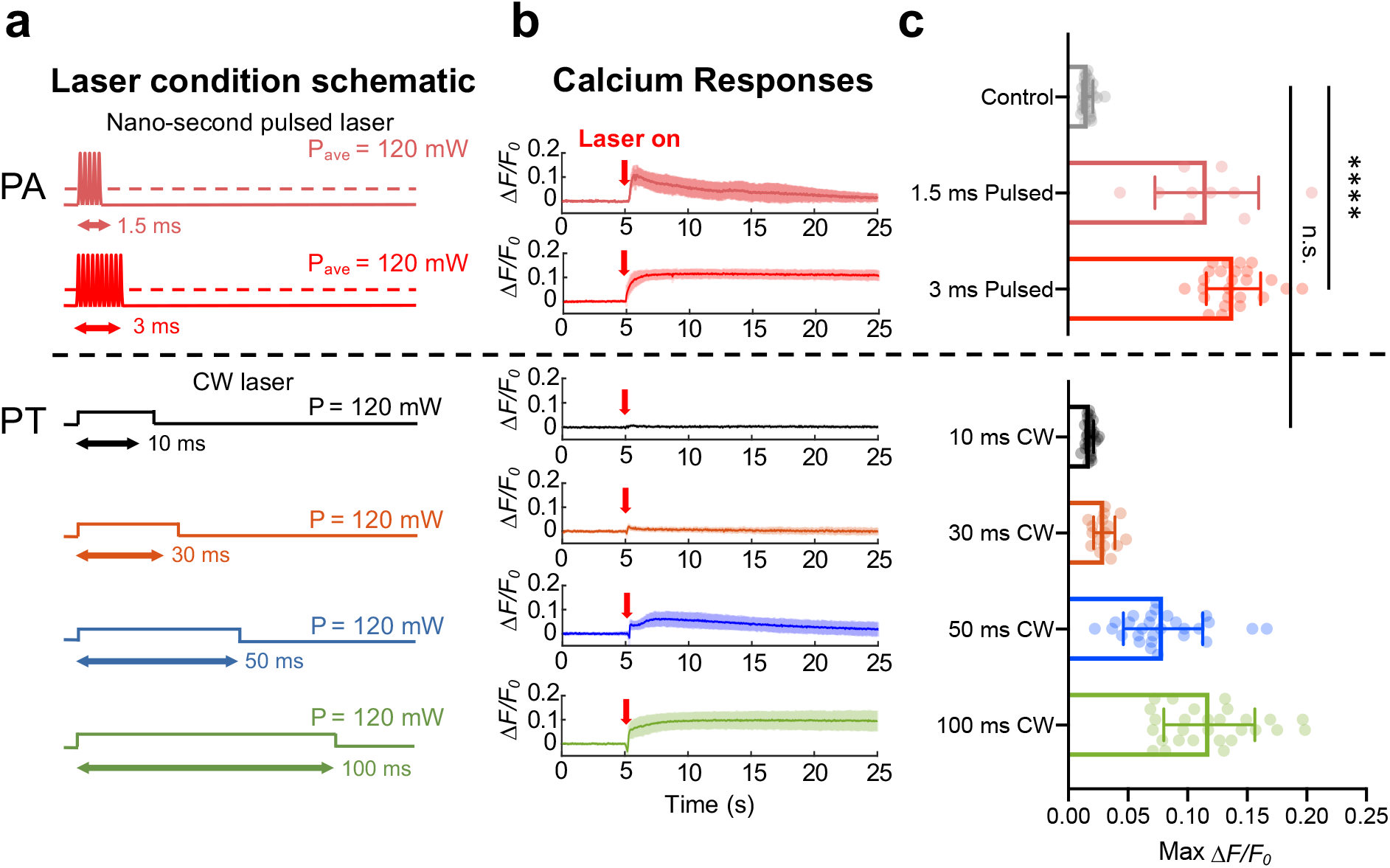
Comparison of neuron response upon PA and PT stimulation recorded by Ca^2+^ imaging. **a**. Schematic of different laser conditions used in PA/PT stimulation. **b**. Ca^2+^ traces of neurons under different laser conditions shown in a. Laser power: 120 mW for CW laser and also 120 mW (averaged power) for pulsed laser. Laser on at t = 5 s (Red arrow). Bold line: averaged trace. Shaded area: SD. **c**. Statistical analysis of maximum *ΔF/F*_0_ of neurons under PA and PT stimulation. Control: no laser. N = 28, 10, 25, 28, 16, 28, 28 for control group, 1.5 ms Pulsed, 3 ms Pulsed, 10 ms CW, 30 ms CW, 50 ms CW, 100 ms CW, respectively. *t-test*, ****, p < 0.0001. n.s. no significance.

We then applied the 1064 nm CW laser to the same FE. The CW laser was operated at the same average power of 120 mW (**Fig. 2a**), acting as a purely PT source. The absorption spectrum of the candle soot under both the 1030 nm and 1064 nm was shown to be the same (**Supplementary Figure S3**). Therefore, the total energy absorbed by FE under these two laser conditions is expected to be the same.

Unlike the pulsed laser condition, employing a CW laser of 10 ms duration with the same average power failed to elicit neuronal stimulation within the area of interest (**Fig. 2b**, black trace). This result shows that even though we delivered nearly three times the laser energy to the FE, the PT stimulation threshold was not reached, and the fluorescence change was minimal (1.8 ± 0.3%). When the CW laser duration increased to 30 ms, a fluorescence response of 3.0 ± 0.9% was observed (**Fig. 2b**, orange trace). Notably, when increasing the CW laser duration to 50 ms, the fluorescence change reached 7.9 ± 3.3% and clear activation of the neurons was observed (**Fig. 2b**, blue trace). Finally, with the CW laser duration reaching 100 ms, the neurons were effectively stimulated (11.8 ± 3.8%), displaying a fluorescence trace akin to that observed under the 3 ms photoacoustic condition (**Fig. 2b**, green trace). We further conducted statistical *t-test* analysis to support our findings (**Fig. 2c**). Specifically, no significant difference between the control group and the 10 ms PT group (n.s., p = 0.172). However, a significant difference was observed between the 3 ms PA group and the 10 ms PT group (****, p < 0.0001). Based on this comparative analysis, it becomes evident that the heat or temperature increase generated under the 3 ms pulsed laser condition is insufficient to stimulate neurons.

Considering the first 0.6 ms of idle time when the pulsed laser was turned on (**Supplementary Figure S5**), only 2.4 ms was needed for PA stimulation. Thus, compared to the 100 ms laser duration needed for PT stimulation, approximately 1/40 of the energy dosage is required for PA to achieve a similar level of neuron activation.

### Patch clamp recording shows distinct neuronal response to PA and PT stimulations

While Ca^2+^ imaging showed distinct energy dose requirements for PA and PT stimulation, it is an indirect measurement of neuronal activities. Instead, patch clamp recording can provide direct recording of sub- and supra-threshold neuron activities. Previously, patch clamp recording of ultrasound stimulation has been limited as conventional transducer-generated ultrasound easily disrupts the patch attachment. One unique capability of our device is its compatibility with whole-cell patch-clamp recording of single neurons ^24^. Here, we evaluated FE-invoked electrical responses in single cultured cortical neurons with whole cell patch clamp (**Fig. 3a**). In order to be compatible with the patch-clamp recording system, we used a tapered FE with a tip diameter smaller than 50 µm (**Fig. 3b**) ^24^. It can generate an ultrasound field confined to about 80 µm and enabled single-neuron stimulation in a low-density neuron culture (< 20 cells/mm^2^).

**Figure 3.**
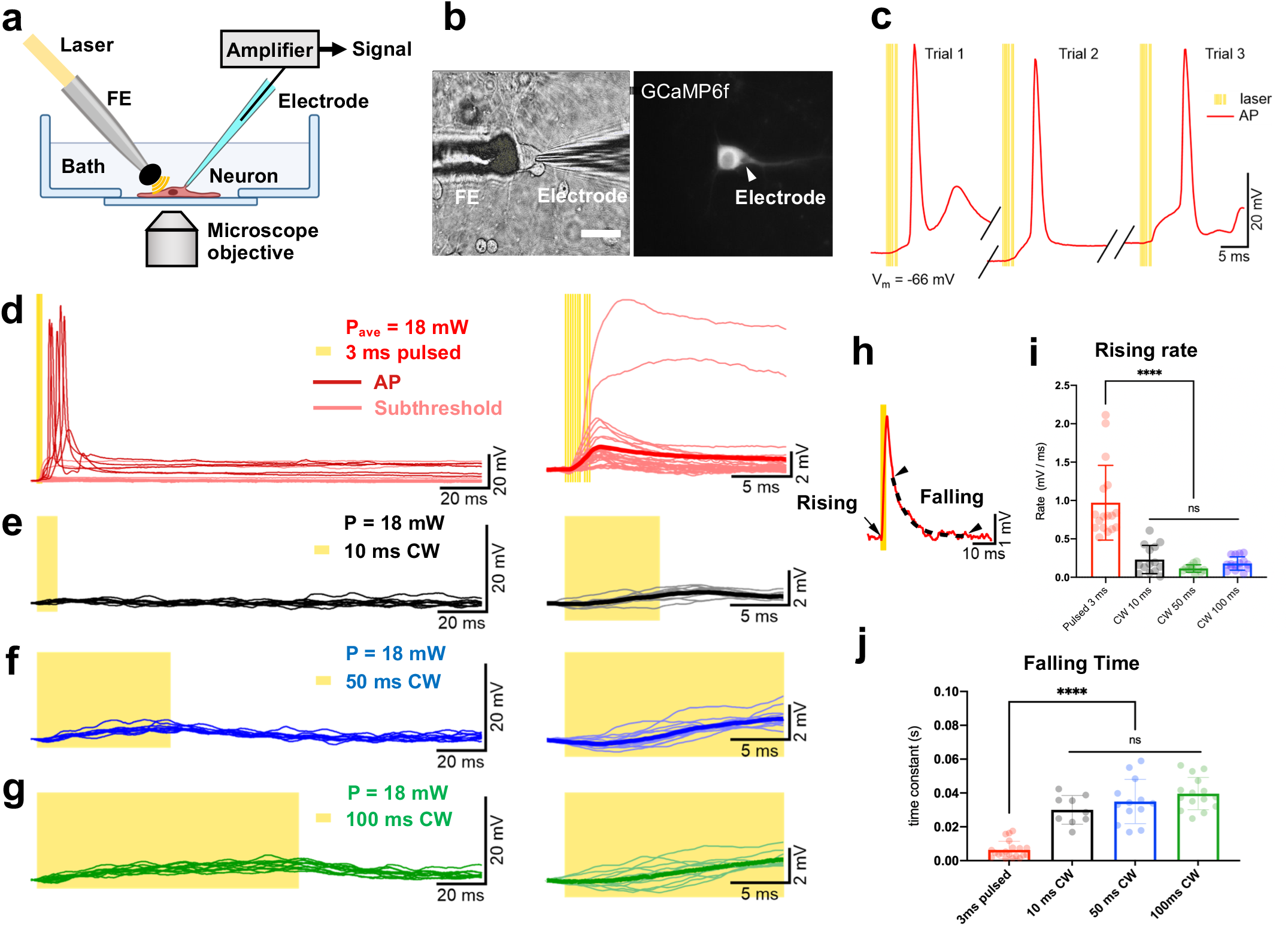
PA-triggered subthreshold depolarizations and action potentials (APs) are distinct from PT effects. **a**. Schematic of tapered FE stimulation and electrophysiological recording. **b**. Transmission and fluorescence images of simultaneous whole cell patch clamp and PA stimulation on the same neuron. Left: transmission light. Right: GCaMP6f fluorescence imaging. Scale bar: 50 µm. **c**. Representative APs triggered by FE stimulation in 3 consecutive trials on one neuron. Each yellow line denotes a 3 ns laser pulse. **d**. Overlaying representative single trial recordings of membrane responses, including APs and subthreshold membrane depolarizations, induced by 3 ms pulsed laser (N=41, 6 cells). Dark red traces: APs. Pink: subthreshold depolarizations. Right panel: the same traces, excluding APs (N=27, 4 cells), at the early stage of the depolarizations. Bold red line: average of all the subthreshold depolarizations. **e-g**. Overlaying representative single traces of membrane responses triggered by CW laser of 10, 50, and 100 ms duration (N=17, 3 cells). Yellow shaded area denotes laser on. Right panel: the same traces showing the beginning of the depolarizations. Bold line: average of the traces in each condition. All traces in d-g were normalized to the same baseline 2 ms before laser delivery. **h**. An example trace from panel d indicating the selected time points for measurements in i and j. The black arrow when the rising rate (black arrow) in i was measurement. Two arrow heads points out the start and end of the selected period for calculating time constant in the decay phase in j. Dashed line: single exponential fit of the selected curve between the arrow heads. **i**. Statistical summary of the rising rate of the subthreshold depolarizations at the very beginning in respond to FOE stimulation. *ANOVA* test. ****, p<0.0001. **j**. Statistical comparison of the time constant of the decay phase in PA and PT-triggered subthreshold depolarizations. *ANOVA* test. ****, p<0.0001. APs in d were excluded from statistical analysis in i and j for comparison of only subthreshold events.

The nanosecond pulsed laser at 1030 nm with a 4.23 kHz repetition rate or the CW laser at 1064 nm was delivered to the tapered FE. The averaged power of each type of laser delivered at the tapered tip was measured to be 18 mW (N=10). As shown in **Fig. 3a** (red line), the FE with a ns pulse train of 3 ms duration could directly trigger action potential (AP) firing in consecutive trials on single neurons (**Fig. 3c**). Membrane started to depolarize immediately following the first pulse, gradually preceded with a polynomial curve until reaching the AP threshold with an average delay of 3.99 ± 3.29 ms after laser onset (**Fig. 3c, 3d**). Among the neurons tested (N=19 from 5 animals), 58% responded to FE-based PA stimulation. Among those responded to FE stimulation, 36% responded with the occurrence of AP, presumably due to a lower laser energy dosage compared to the previous calcium imaging experiments and resulted in a stimulation power around or below the threshold of AP firing.

Additionally, the characteristics of the PA triggered APs are found similar to that of APs occurred spontaneously (**Supplementary Figure S4 a-c**), including maximum amplitude, after-hyperpolarization, AP half-width, rising rate, and falling rate (**Supplementary Figure S4 d-f**). In most other responding cases, the same pulsed laser repeatedly triggered subthreshold depolarizations with an average amplitude of 1.84 ± 0.98 mV (**Fig. 3d**, pink curves). The largest triggered amplitude was 13.57 mV while the smallest was 0.16 mV. The variations were likely caused by differential cell sensitivity to PA stimulation. The maximum amplitude occurred at 0.36 ± 0.66 ms after the laser offset on average. These data suggest that like spontaneous firing, ion channels are likely involved in PA stimulation.

In contrast to the PA condition, no APs were triggered by the FE driven by the CW laser of the same power of 18 mW, for durations up to 100 ms. The 10 ms CW laser induced membrane depolarizations with a maximum amplitude of 0.93 ± 0.40 mV at 2.90 ± 0.19 ms after laser offset (**Fig. 3e**). Membranes of the same neurons were depolarized for 2.57 ± 0.24 mV with 50 ms CW laser (**Fig. 3f**) and 5.24 ± 0.27 mV with 100 ms CW laser (**Fig. 3g**). The depolarizations reached a maximum at 2.86 ± 8.55 ms before and 1.55 ± 4.78 ms after laser offset in the 50 ms and 100 ms conditions, respectively. At 3 ms after laser onset in all CW laser conditions, the averaged membrane response was −0.042 ± 0.50 mV, almost the same as the baseline, showing that the 3 ms of heating is not enough to induce obvious membrane depolarization.

Besides the difference in amplitudes and variations, distinct characteristics were shown in the rising and falling phases of the membrane responses to the PA and PT stimulations. Membrane depolarizations triggered by the pulsed ns laser showed a much faster responding and decay rate compared to those by CW laser. To better compare the properties of the subthreshold events, we analyzed the rising rate at the beginning of the responses, and the time constant of the recovering periods by a single exponential fit (**Fig. 3h**) in each laser condition. In the pulsed laser condition, subthreshold events had an average rising rate of 0.97 ± 0.49 mV/ms. In 10 ms, 50 ms, and 100 ms CW laser condition, the rising rates were 0.23 ± 0.18 mV/ms, 0.11 ± 0.049 mV/ms, and 0.18 ± 0.086 mV/ms, respectively. ANOVA test showed a significantly larger membrane depolarization speed in the PA condition compared to all of the PT conditions (****, p<0.0001, **Fig. 3i**).

At the decay phase, a time constant of 6.34 ± 5.09 ms suggested a much faster decay phase of PA-triggered subthreshold depolarizations compared to the PT triggered ones (30.02 ± 8.49 ms for 10ms, 34.98 ± 13.1 ms for 50ms, 39.65 ± 9.56 ms for 100 ms). Notably, membrane responses in PA conditions demonstrated a two-phase decay, instead of a single exponential component, which indicates the activation of active membrane components, namely the opening of ion channels^24^, in addition to passive responses of the lipid membrane. Accordingly, statistical summary suggests a significant difference in the decay time between the PA and PT-triggered subthreshold depolarizations (****, p<0.0001, **Fig. 3j**). No significant difference was found within PT conditions in the rising rate (n.s., p = 0.076, **Fig. 3i**) or the time constant (n.s., p = 0.115, **Fig. 3j**), indicating they all resulted from temperature-induced capacitance responses on the cell membrane.

Collectively, these results show that PA and PT have triggered distinct electrical responses. In consistence with the calcium imaging data, our patch data support that thermal effect did not play a major role in the membrane responses to the pulsed laser and that PA should trigger the membrane depolarizations through a non-thermal mechanism.

### Blocking mechanosensitive ion channels effectively suppresses PA stimulation

The distinct membrane responses to PA versus PT stimulation inspired us to investigate the molecular mechanisms. Previous studies have suggested that mechanosensitive ion channels play a significant role in ultrasound neurostimulation ^3^. Thus, we hypothesized that PA alters the mechanical properties of the plasma membrane and subsequently induces activation of mechanosensitive channels. These channels accumulate ionic currents to form membrane depolarization, and subsequently activate voltage-gated sodium channels to induce action potentials. To test this hypothesis, the contribution of mechanosensitive channels and voltage-sensitive sodium channels in neuromodulation was assessed by pharmacological inhibition of channel activity.

To compare the PA stimulation results with blockers of different ion channels, a control experiment on GCaMP6f-neurons without any pharmacological treatment was performed. Syn-driven GCaMP6f, a Ca^2+^ sensor, was delivered to neurons via AAV9 viral vector at 4 days in vitro (See Methods). By applying a 3 ms pulse train of ns pulses with an energy of 24 µJ per pulse to an FE of 100 µm in diameter, the PA signal generated successfully stimulated the surrounding neurons with an averaged Max Δ*F*/*F*_0_ = 122.1 ± 75.2%. This result is considered as the baseline for other neuron groups with different kinds of blockers under the same PA stimulation.

First, to evaluate the involvement of thermosensitive ion channels, Ruthenium red was applied to the neurons to block TRPV1, TRPV2 and TRPV4 channels ^25, 26^. As shown in **Fig. 4a, 4g**, no significant difference was observed compared to the control group (n.s., p = 0.28), indicating that these channels were not involved under this PA condition. Similar findings were also reported in transducer-based ultrasound stimulation ^3^.

**Figure 4.**
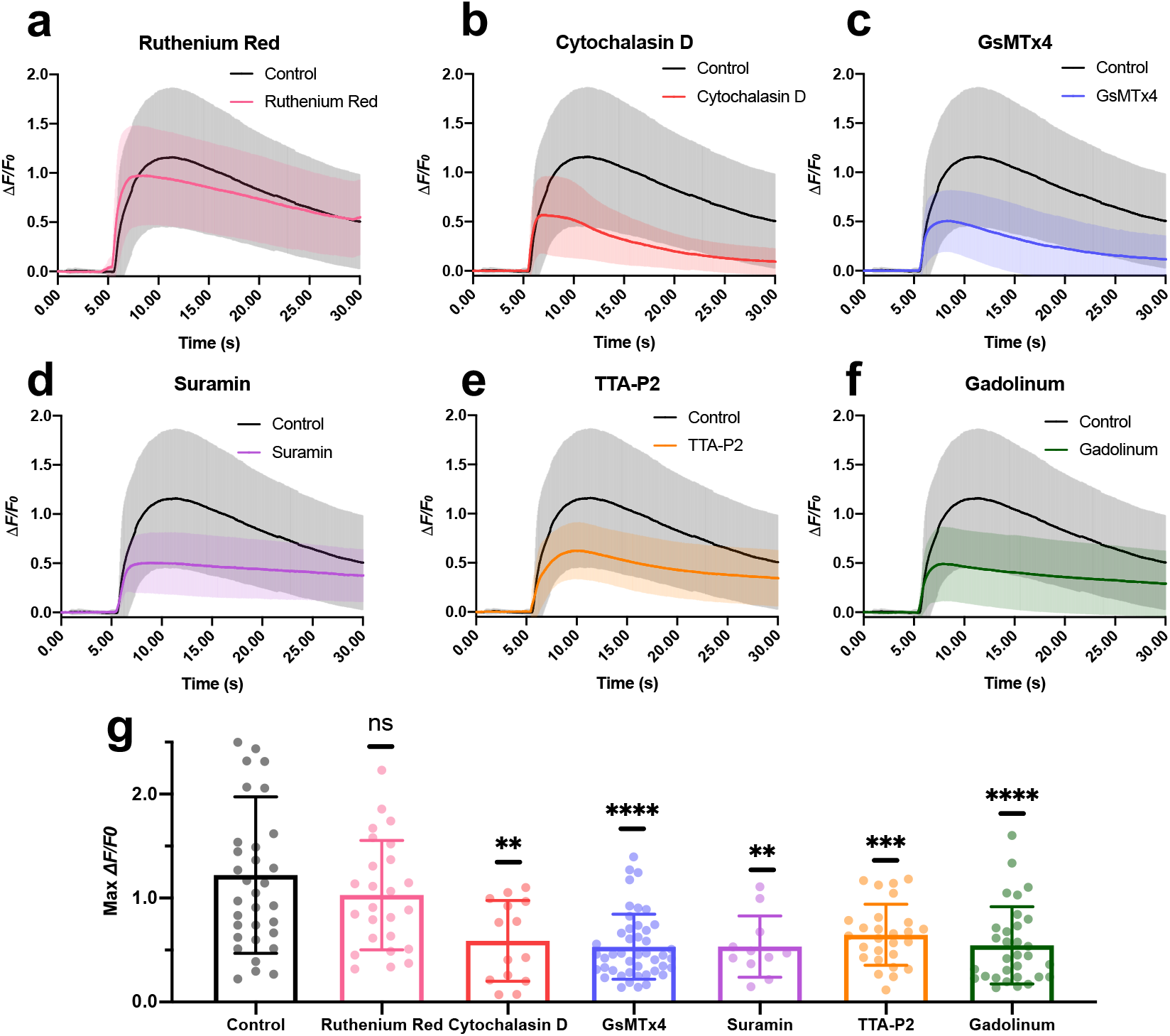
Bocking mechanosensitive ion channels effect PA stimulation of cortical neurons. **a-f**. Calcium traces of neurons treated with blockers (Ruthenium Red: pink, N = 25. Cytochalasin D: red, N = 14. GsMTx4: blue, N = 45. Suramin: purple, N = 11. TTA-P2: orange, N = 27. Gadolinum: green, N = 31. Control: Black, N = 33.) upon FE stimulation. Shaded area: SD. **g**. Statistics of the max *ΔF/F*_*0*_ under varied treatments. *t-test* n.s. not significant. ****, p < 0.0001. ***, p < 0.001. **, p < 0.01.

Next, to alter the mechanical properties of the neurons, cytochalasin D was added to the neuron culture to inhibit the membrane ruffling by depolymerizing the actin cytoskeleton ^27^. The Max *ΔF/F*_*0*_ decreased by 63.3% (**, p = 0.0047) (**Fig 4b, 4g**), indicating that the elastic modulus is important during the acoustic-induced membrane distortion. Next, the peptide inhibitor GsMTx4, which blocks Piezo1 and TRPC1 channels, was used. The Max *ΔF/F*_*0*_ of neurons showed a decrease of 69% in response to the FE-based PA stimulation (****, p < 0.0001) (**Fig 4c, 4g**), indicating that the Piezo1 and/or TRPC1 channels play a key role during this process.

G protein-coupled receptors (GPCRs) are sensory molecules reported to be important for mechano-transduction in vasculature as shear stress sensors ^28^. To investigate whether the GPCRs were activated, suramin was used to block GPCR signaling (**Fig. 4d, 4g**). The Max *ΔF/F*_*0*_ decreased by 68.9% (**, p=0.0053), suggesting the shear stress promoted the signaling of GPCRs for cell activation. It is also worth noting that, in the recent work using focused ultrasound for neurostimulation, GPCRs were shown to be not involved in the stimulation process ^3^. This discrepancy might originate from the difference in acoustic wave propagation. In the focused-ultrasound study, the spherical focal area of the acoustic wave with a 5 mm diameter could cover the entire cell culture and be regarded as a planar wave. While in our FE work, the generated acoustic field has a sub-millimeter diameter and propagates omnidirectionally, denoting a point source. Thus, shear stress was likely to be present in the lateral wave propagation of the FE-generated ultrasound field.

Besides, L-type, N-type, T-type, and P-type calcium channels have been shown to be mechanically sensitive under various conditions ^29, 30^. Recently, voltage-gated T-type calcium channels was reported to be downstream amplifiers for ultrasound neuromodulation ^3^. To validate this, we treated the cells with the selective blocker TTA-P2, which suppressed the Max *ΔF/F*_*0*_ by 57.4% (***, p = 0.0004) (**Fig. 4e, 4g**). Thus, voltage-gated T-type calcium channels are likely activated during PA stimulation.

Lastly, we tested the effect of Gadolinium(III), which has been reported as a nonspecific agent that blocks mechano-gated channels via changing the deformability of the lipid bilayer ^31^. As shown in **Fig. 4f**, the FE-induced calcium activities were significantly inhibited. Statistical results (**Fig. 4g**) show that Gadolinium(III) resulted in a *ΔF/F*_*0*_ decrease by 67.5% (****, p<0.0001). These data collectively demonstrate the involvement of mechanosensitive ion channels in PA stimulation.

### Overexpressing TRPC1/TRPP2/TRPM4 boosts the calcium signal upon PA stimulation

According to an earlier report that ultrasound excites neurons via the activation of endogenous mechanosensitive ion channels, including TRPP2 and TRPC1, as well as the calcium-dependent amplifier TRPM4 channel ^3^, we overexpressed these three ion channels to identify their roles in PA stimulation. The gene constructs for TRPC1, TRPP2, and TRPM4 ion channels were overexpressed in GCaMP6f neurons under a hSyn promoter (**Fig. 5a**). The fluorescent protein mCherry was co-expressed as an expression indicator (**Fig. 5b**). To further quantify the overexpression of ion channels, immunofluorescent labeling was performed in the control group and the overexpression groups. As shown in **Fig. 5c-e**, the signals for TRPC1, TRPP2 and TRPM4 channels in the overexpression groups are 33 ± 8%, 30 ± 15%, 32 ± 14% higher than the wild type groups, respectively, suggesting successful and comparable overexpression.

**Figure 5.**
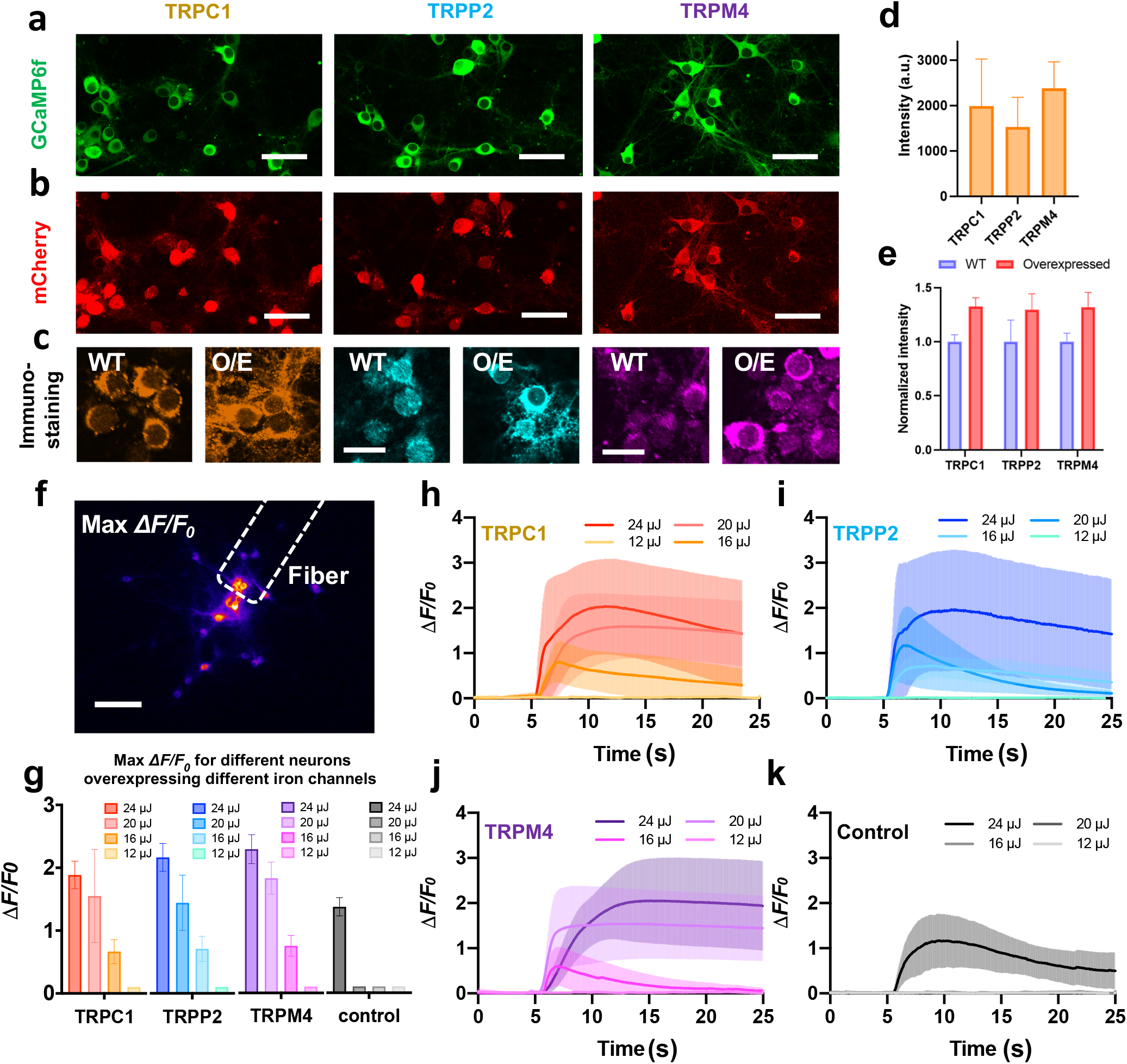
Response of neurons to PA stimulation after the overexpression of mechanical sensitive ion channels. **a**. Neurons expressing GCaMP6f as calcium indicator. **b**. Co-expressing of mCherry with the TRPC1/TRPP2/TRPM4 ion channels. Scale bars: 50 µm. Right panel: statistics of the mCherry signal intensity. **c**. Immunostaining of ion channels in wild type (WT) group and overexpression (O/E) group. Scale bars: 20 µm. **e**. Statistics of the immunostaining signal intensity. **f**. Representative max *ΔF/F*_*0*_ contrast imaging of tapered FE based PA stimulation. Dashed line: location of the tapered FE. Scale bar: 100 µm. **g**. Analysis of the calcium responses of neurons overexpressing ion channels upon FE stimulation with varied laser pulse energy. Error bars: SD. **h-k**. Calcium traces of neurons overexpressing TRPC1/TRPP2/TRPM4 channels and control group upon FE stimulation with varied laser pulse energy. SD.

Next, the FE-based PA stimulation was performed on neurons overexpressing specific ion channels. A 3 ns pulsed laser at 1030 nm and a 1.7 kHz repetition rate was used to deliver laser pulses of 3 ms duration to a FE with a 100 μm tip diameter (**Fig. 5f**). Varied laser pulse energy of 12, 16, 20, and 24 µJ were applied to test the neuron activation threshold (**Fig. 5h-5k**). As shown in **Fig. 5h-5i**, the PA stimulation of neurons overexpressing TRPC1 and TRPP2 evoked substantially larger calcium activities (TRPC1: Max *ΔF/F*_*0*_ = 188.5 ± 21.9%, TRPP2: Max *ΔF/F*_*0*_ = 216.7 ± 22.2%) compared to the control group (Max *ΔF/F*_*0*_ = 136.5 ± 14.4%) with pulse energy of 24 µJ. Meanwhile, under the pulse energy of 16 or 20 µJ, the TRPC1 and TRPP2 groups could be stimulated (TRPC1: Max *ΔF/F*_*0*_ at 16 µJ = 66.5 ± 19.1%, Max *ΔF/F*_*0*_ at 20 µJ = 155.0 ± 74.0%; TRPP2: Max *ΔF/F*_*0*_ at 16 µJ = 71.0 ± 20.0%, Max *ΔF/F*_*0*_ at 20 µJ = 144.3 ± 44.0%), while no activity was observed in the control group, indicating that the overexpressing TRPC1 and TRPP2 channels increased the mechanosensitive of the neurons (**Fig. 5g**). Likewise, neurons overexpressing the TRPM4 channel showed increased calcium response and a lower stimulation threshold (Max *ΔF/F*_*0*_ at 24 µJ = 228.0 ± 23.0%, Max *ΔF/F*_*0*_ at 20 µJ = 182.0 ± 25.3%, Max *ΔF/F*_*0*_ at 16 µJ = 74.8 ± 16.5%), validating the involvement of TRPM4. It is conceivable that TRPM4 serves as the downstream calcium-dependent amplifier even though itself is not mechanosensitive (**Fig. 5j**). Collectively, these data demonstrate the involvement of TRPC1, TRPP2, and TRPM4 channels in PA stimulation.

### The optocapacitive mechanism is not involved in PA stimulation

In addition to the amount of temperature elevation, an optocapacitive mechanism through rapid temperature increase was proposed for PT stimulation ^11, 32^. To test the possible involvement of the optocapacitive mechanism in PA stimulation, we harnessed an 80-MHz femtosecond (fs) laser to FE to deliver a rapid temperature increase in 3 ms period.

The Newton’s law of heating states that mC_s_(Δ*T*) = *Q*_*abs*_, where m is the mass and Cs is the heat capacity. In the same period of 3 ms, if the total heat *Q*_*abs*_ is the same, Δ*T*, the total temperature rise should be the same over the same period. For each pulse, the temperature rising slope is proportional to the heat transfer rate, which is linear to the pulse peak power. In our case, we used a nanosecond laser with 4.23 kHz repetition rate, 3 ns pulse duration, 120 mW averaged power, and a fs laser with 80 MHz repetition rate, 140 fs pulse duration with the same averaged power of 120 mW. According to P_peak_=P_ave_/(R_f_*τ), where R_f_ and τ are laser repetition frequency and pulse duration, respectively, the peak power of the fs laser condition is 1.13 times that of the ns laser. Based on this rationale, we expect that the fs laser delivers a similar temperature increase rate as the ns laser does and can be used to evaluate the optocapacitive contribution in the PA stimulation. With the same peak power and averaged power, the fs laser has a large repetition rate of 80 MHz, corresponding to a small pulse energy (1.5 nJ/pulse). Since the pressure from the photoacoustic effect is related to the pulse energy, the PA effect from this fs laser condition is negligible.

Experimentally, under the fs laser condition, with the same burst duration of 3 ms, the FE failed to stimulate neurons (**Supplementary Figure S6, black line**, N = 24). In contrast, under the same burst duration, the ns laser can evoke neurons efficiently with a Max *ΔF/F*_*0*_ larger than 5% (**Supplementary Figure S6, red line**, N = 10). This result indicates that the rapid temperature increase that occurred during the fs laser condition is insufficient to stimulate neurons. Under the ns laser condition, despite the presence of the same temperature increase rate, it is the PA effect that successfully stimulated the neurons.

## Discussion

In this work, using a unified fiber emitter of heat and ultrasound, we compared the laser energy needed for PA and PT stimulation. The laser energy needed for PA stimulation was shown to be 1/40 of that for PT stimulation. This substantial difference demonstrates that the PT effect associated with the PA process alone is not sufficient to trigger neurons; instead, the generated ultrasound is the key factor for neurostimulation. Practically, the lower energy required for PA stimulation is advantageous in two perspectives. First, it reduces the potential risk of thermal damage in neural tissues. Second, the lower energy demand opens up the potential of applying PA stimulation in deeper tissue as it can afford more energy loss due to scattering.

The whole-cell patch configuration has long served as the golden standard for studying neuronal responses to stimuli. However, measuring cellular responses under transducer-produced ultrasound has proven exceptionally challenging due to the widespread interference of ultrasound with the patch, and only a few studies ^33, 34^ have reported measurements under pressures up to 200 kPa. This study demonstrates that whole-cell patch measurements of individual cultured neurons are possible under high-precision PA and PT stimulation. Our findings showed that PA-induced membrane depolarizations had significantly faster rising and falling rates, indicating the involvement of ion channel activation. In contrast, the depolarization amplitudes in PT stimulation were sensitive to temperature change, with the rising and falling rate of depolarizations aligned with the rate of temperature change. These results suggest a predominant role of a heat-induced capacitance mechanism ^10, 11^ in PT stimulation, rather than the activation of temperature-sensitive ion channels. Collectively, these findings highlight distinct molecular mechanisms underlying the depolarization responses to PA and PT stimulations.

We further investigated the molecular mechanisms underlying PA modulation. Since the PA devices generate acoustic waves in the ultrasonic range, it is conceivable that PA neurostimulation shares similar mechanisms with ultrasound neurostimulation. To date, several mechanisms for ultrasound stimulation have been proposed, including local temperature increase ^35^, transient sonoporation ^36, 37^, intramembrane cavitation ^38, 39^, and activation of mechanosensitive ion channels ^3, 40, 41^. Among these possible mechanisms, activation of mechanosensitive ion channels has been the most extensively studied hypothesis for acoustic neuromodulation. In an oocyte membrane system, Kubanek et al. recorded transmembrane currents from individually expressed mechanosensitive ion channels including TREK-1, TREK-2, TRAAK, and Na_v_1.5 ^40^. A later study by Kubanek et al. further identified MEC-4, an ion channel for a touch sensation, was crucial for ultrasound-modulated responses in Caenorhabditis elegans ^41^. In addition, the overexpression of TRP-4, a TRPN family channel has been shown to enhance ultrasound modulation in Caenorhabditis elegans as well ^42^. Employing calcium imaging, Gaub et al. investigated the neuronal response to pure mechanical stimuli using atomic force microscope cantilever ^43^. They identified the force and pressure required for both transient and sustained activation. The contribution of various mechanosensitive ion channels has also been explored through pharmacological manipulation. Using calcium imaging, Yoo et al. examined the activation of various mechanosensitive ion channels upon ultrasound stimulation and identified the key contribution of three ion channels: TRPP2, TRPC1 and Piezo1 ^3^. The proposed downstream molecular pathway involves calcium amplification by TRPM4 and voltage gated calcium channels. In this work, we showed PA stimulation of primary cortical neurons through specific calcium-selective mechanosensitive ion channels with the assistance of calcium amplifier channel and voltage-gated channels. As our key findings, pharmacological inhibition of specific ion channels leads to reduced responses, while over-expressing TRPC1, TRPP2 and TRPM4 channels results in stronger stimulation. Collectively, these results shed new insights into the mechanism of PA and ultrasound neurostimulation.

## Materials and Methods

### FE fabrication and characterization

For Ca^2+^ imaging of PA and PT neuromodulation, we used a multimode fiber (FT200EMT, Thorlabs, Inc., NJ, USA) with 200 µm in diameter. The PA coating was composed of candle soot and PDMS. Candle soot was chosen as the absorber due to its great absorption coefficient. The multimode optical fiber was exposed the candle flame for around 3-5 seconds until the fiber tip was fully coated, with the thickness of candle soot controlled by the deposition time ^20^. To prepare PDMS, the silicone elastomer (Sylgard 184, Dow Corning Corporation, USA), was carefully dispensed into a container to minimize air entrapment, and then mixed with the curing agent in a ratio of 10:1 by weight. A nanoinjector deposited the prepared PDMS onto the tip of the candle-soot coated fiber and thus formed a layered structure ^44^. The position of the fiber and the nanoinjector were both controlled by 3D manipulators for precise alignment, and the PDMS coating process was monitored in real-time under a lab-built microscope. The coated fiber was then cured overnight at room temperature.

For stimulation performed with whole cell patch with single cell stimulation, tapered FE was prepared using multimode optical fibers was scribed (S90C, Thorlabs, Inc., NJ, USA) as previously described ^24^. For pharmacological blocking and ion channel studies, the same type of tapered FE was used.

To characterize the photoacoustic signal generated by the FE, a customized and compact passively Q-switched diode-pumped solid-state laser (1030 nm, 3 ns, 100 µJ, repetition rate up to 10 kHz kHz, RPMC, Fallon, MO, USA) served as the excitation source. The laser was first connected to an optical fiber through a customized fiber jumper (SMA-to-SC/PC, ∼81% coupling efficiency) and then connected to the FE with a SubMiniature version A (SMA) connector. To adjust the laser power, fiber optic attenuator sets (multimode, varied gap of 2/4/8/14/26/50 mm, SMA Connector, Thorlabs, Inc., NJ, USA) were used. A needle hydrophone (ID. 40 µm; OD, 300 µm) with a frequency range of 1-30MHz (NH0040, Precision Acoustics Inc., Dorchester, UK) was utilized for the acoustic measurement. Both the fiber and the hydrophone were aligned under the microscope (**Supplementary Figure S1**). The acquired signal was processed with an ultrasonic pre-amplifier (0.2–40MHz, 40 dB gain, Model 5678, Olympus, USA) and a digital oscilloscope (DSO6014A, Agilent Technologies, USA). The distance between the FE tip and hydrophone was controlled using a 4-axis micro-manipulator (MC1000e controller with MX7600R motorized manipulator, Siskiyou Corporation, USA) with a controllable motion of 0.2 µm. The distance was measured using a widefield microscope with a 20× objective. Both the FE tip and hydrophone tip were immersed in degassed water dropped on a cover glass. The pressure values were calculated based on the calibration curve obtained from the hydrophone manufacturer. The frequency data was obtained through the Fast Fourier Transform (FFT) using MATLAB 2020a.

### Neuron culture with GCaMP6f / mCherry expression

The glass-bottom culture dishes used in the embryonic neuron cultures were immersed in 0.01% Poly-D-Lysine (Sigma-Aldrich, MO) overnight at 4 °C and washed in PBS twice before culture initiation. Primary cortical neurons were obtained from SD rat E15 embryos (Charles River Laboratory). Dissociated cells were washed and triturated with 10% heat-inactivated fetal bovine serum (FBS, Atlanta Biologicals, GA), 5% heat-inactivated horse serum (HS, Atlanta Biologicals, GA), 2 mM Glutamine-Dulbecco’s Modified Eagle Medium (DMEM, Thermo Fisher Scientific Inc., MA), and cultured in cell culture dishes (100 mm diameter) for 30 min at 37 °C to eliminate glial cells and fibroblasts. The supernatant containing neurons was collected and seeded on poly-D-lysine coated cover glass and incubated in a humidified atmosphere containing 5% CO_2_ at 37 °C with 10% FBS + 5% HS + 2 mM glutamine DMEM. After 16 h, the medium was replaced with Neurobasal medium (Thermo Fisher Scientific Inc., MA) supplemented with 2% B27 (Thermo Fisher Scientific Inc., MA), 1% N2 (Thermo Fisher Scientific Inc., MA), and 2 mM glutamine (Thermo Fisher Scientific Inc., MA). Half of the medium was changed with the fresh medium every 3 days, and neurons were used for stimulation experiments after 12–14 days from the seeding.

For calcium imaging, Syn-driven GCaMP6f as a calcium sensor was delivered to neurons via AAV9 viral vector transfection (Addgene, pAAV.Syn.GCaMP6f.WPRE.SV40, 1E10 vp/dish) at 4 days in vitro. To measure temperature change during photoacoustic and photothermal stimulation, mCherry as a temperature sensor was delivered to neurons via AAV9 viral vector transfection (Addgene, pAAV-hSyn-mCherry, 1E10 vp/dish). The viral particles were added to neurons at 3 days in vitro. The whole media was replaced with the fresh media at 4 days in vitro, and the cells were maintained for 6 additional days.

### Calcium imaging of FE induced neuron stimulation

Calcium fluorescence imaging was performed on a lab-built wide-field fluorescence microscope. The microscope was based on an Olympus IX71 microscope frame, with a 20X air objective (UPLSAPO20X, 0.75NA, Olympus), illuminated by a 470 nm LED (M470L2, Thorlabs) and a dichroic mirror (DMLP505R, Thorlabs). A 3-D micromanipulator (Thorlabs, Inc., NJ, USA) positioned the FE at an angle of 45° to the cells, maintaining a distance of approximately 50 µm between the FE tip and the culture. Image sequences were acquired with a scientific CMOS camera (Zyla 5.5, Andor) at 20 frames per second. The fluorescence intensity analysis was performed using ImageJ (Fiji).

In vitro PA neurostimulation experiments were performed using a Q-switched 1,030-nm nanosecond laser (Bright Solution, Inc. Calgary Alberta, CA). In vitro PT neurostimulation experiments were performed using a CW diode pumped laser (Cobolt Rumba 05-01 series, Sweden)

### Temperature measurement in vitro using mCherry fluorescence

Fluorescence imaging of mCherry was conducted on a lab-built wide-field fluorescence microscope (the same system used for calcium imaging). With 20X air objective (UPLSAPO20X, 0.75NA, Olympus), neurons were illuminated by a white LED with an mCherry filter cube (562/40 excitation, 641/75 emission and a dichroic mirror, MDF-MCHC, Thorlabs). FE was precisely positioned using the 3-D micromanipulator (Thorlabs, Inc., NJ, USA). Image sequences were acquired with a scientific CMOS camera (Zyla 5.5, Andor) at 20 frames per second. The fluorescence intensity analysis was performed using ImageJ (Fiji).

### Gene overexpression of ion channels in cultured neurons

As described in the previous work ^3^, the mouse TRPV1 (GenBank: AB040873.1), TRPP2 (GenBank: BC053058), TRPM4 (GenBank: BC096475), and human TRPC1 (GenBank: Z73903.1) genes were synthesized commercially (Integrated DNA Technologies) and cloned upstream of an internal ribosome entry site (IRES2) and mScarlet (TRPC1, TRPP2) or mRuby3 (TRPV1, TRPM4) gene. The construct was inserted into the lenti-backbone. The viral particles were added to neurons at 3 days in vitro (1E9 vp/sample) and maintained for 10 days. hSyn-driven mCherry was inserted into the lenti-backbone by Gibson assembly to confirm the gene expression. The viral particles were added to neurons at 3 days in vitro (1E9 vp/sample), whole media was replaced with the fresh supplemented Neurobasal media at 4 days in vitro, and the cells were maintained for 10 additional days.

### Immunostaining characterization of ion channel expression levels

For immunostaining, primary neurons were fixed using ice-cold paraformaldehyde (4% in PBS, VWR) for 10 min at 4 °C, and washed with PBS twice. Nonspecific biding was blocked by 6% bovine serum albumin (Sigma) for 30 min at room temperature, and cells were washed in PBS. Primary antibody anti-TRPC1 (1:200, Alomone Labs), anti-TRPM4 (1:200, Alomone Labs) and anti-TRPP2 (1:200, Alomone Labs) were diluted in 1.5% bovine serum albumin, and incubated with cells at 4 °C overnight. After washing with PBS for 3 times, secondary antibodies (Alexa Fluor 488 (1:500, Invitrogen) diluted in 1.5% BSA were added to neurons for 1 h at 37 °C. Cells were washed with PBS, and imaged using a confocal microscope (FV3000, Olympus).

### Pharmacological treatments with chemical blockers and peptide inhibitors of ion channels

To block voltage-gated sodium channels, tetrodotoxin citrate (ab120055, Abcam, MA, USA) was added to the culture to reach 3 µM final concentration 30 min before Calcium imaging. The following blockers were added to medium and incubated with cells for 4h before stimulating cells with optoacoustic: Actin filaments were depolymerized by their specific inhibitors, cytochalasin D (final conc.: 10 µM). Gadolinium was applied to nonspecifically block the mechanosensitive ion channels (final conc.: 20 µM). Ruthenium red (final conc.: 1 µM) was used before ultrasound stimulation to block TRP channels (TRPV1, 2, 4) and TTA-P2 (final conc.: 3 µM) was added to block T-type calcium channels. To inhibit GPCRs, suramin was added (final conc.: 60 µM). GsMTx4 was added to medium (final conc.: 10 µM) to inhibit Piezo1 and TRPC1 channels.

### Electrophysiology and action potential analysis

Membrane potentials were recorded in current clamp mode with an Axopatch 200B amplifier (Axon Instruments, Union City, CA, USA) and digitized with a NI 6251 board (National Instruments, Austin, TX, USA). Signals were low-pass filtered at 10 kHz and sampled at 5 kHz. Cells were recorded at a holding potential of -70 mV in a bath solution (140 mM NaCl, 3 mM KCl, 1.5 mM MgCl_2_, 2.5 mM CaCl_2_, 11 glucose and 10 HEPES, pH 7.4). Data analysis only included cells with resting membrane potentials between -60 to -70 mV. The bath solution, heated to 30 °C, was circulated in the petri dish during the whole recording period. The recording electrodes were filled with a K^+^-based internal solution (135 mM K^+^-gluconate, 5 mM NaCl, 2 mM MgCl_2_, 10 mM HEPES, 0.6 mM EGTA, 4 mM Mg^2+^-GTP, 0.4 mM Ma^+^-ATP) and had a resistance ranging from 5 to 10 MΩ. Data were analyzed and visualized with IGOR PRO (Wavemetrics, Lake Oswego, OR, USA). The threshold of an AP was defined as the membrane potential when the dv/dt first exceeds 10 V/s.

## Supporting information

This is supplementary figures for the MS

## Acknowledgments

This work is supported by Brain Initiative R01NS109794 to JXC and CY, and ARO W911NF2110132 to CY. The cortical neurons were provided by Hengye Man lab at Boston University. The Plasmids of mechanosensitive channels were provided by Sangjin Yoo in Mikhail Shapiro lab at Caltech.

## Author Contributions

GC, LS, FY, J-XC and CY: drafting and refining the manuscript. GC: conducting of the PA and PT comparison under calcium imaging. LS: conducting of the pharmacological and genetic studies. FY: conducting of the experiment of patch clamp recording. J-XC and CY: critical guidance of the project. ZD, CM, DJ: help with the experiments. All authors have read and approved the manuscript.

