## Supplementary material for "High-precision photoacoustic neural modulation uses a non-thermal mechanism": This is supplementary figures for the MS

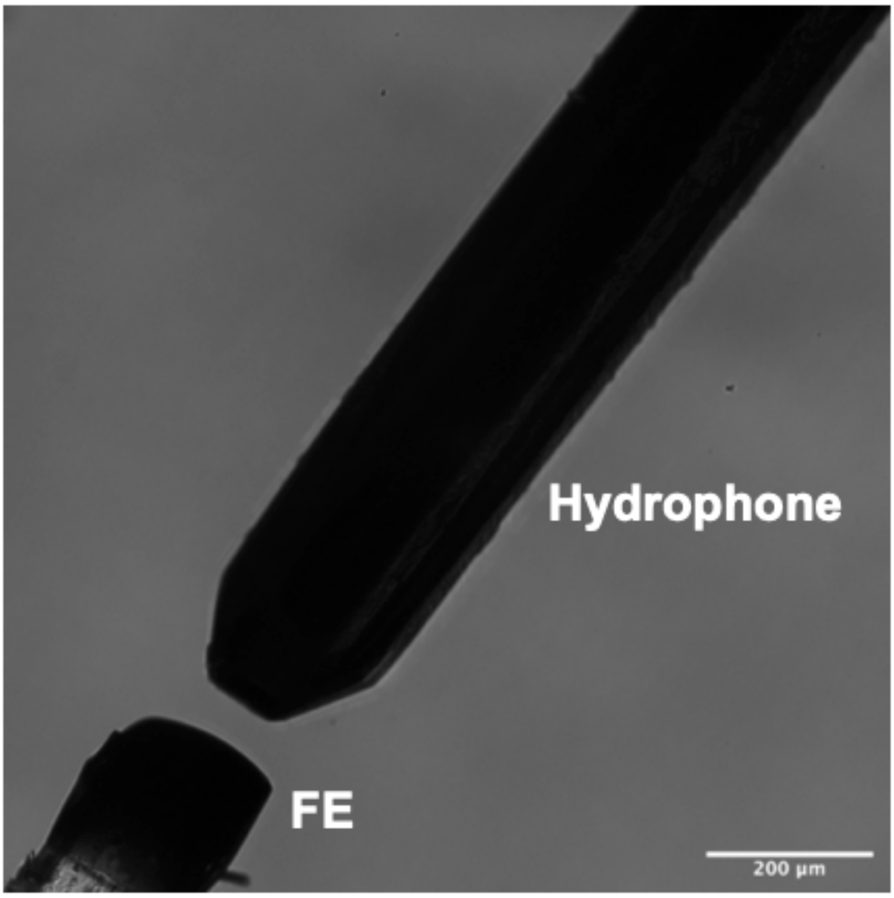


**Figure S1. Measurement of fiber-emitter generated photoacoustic signal by a needle hydrophone.** Left: FE. Right: hydrophone.

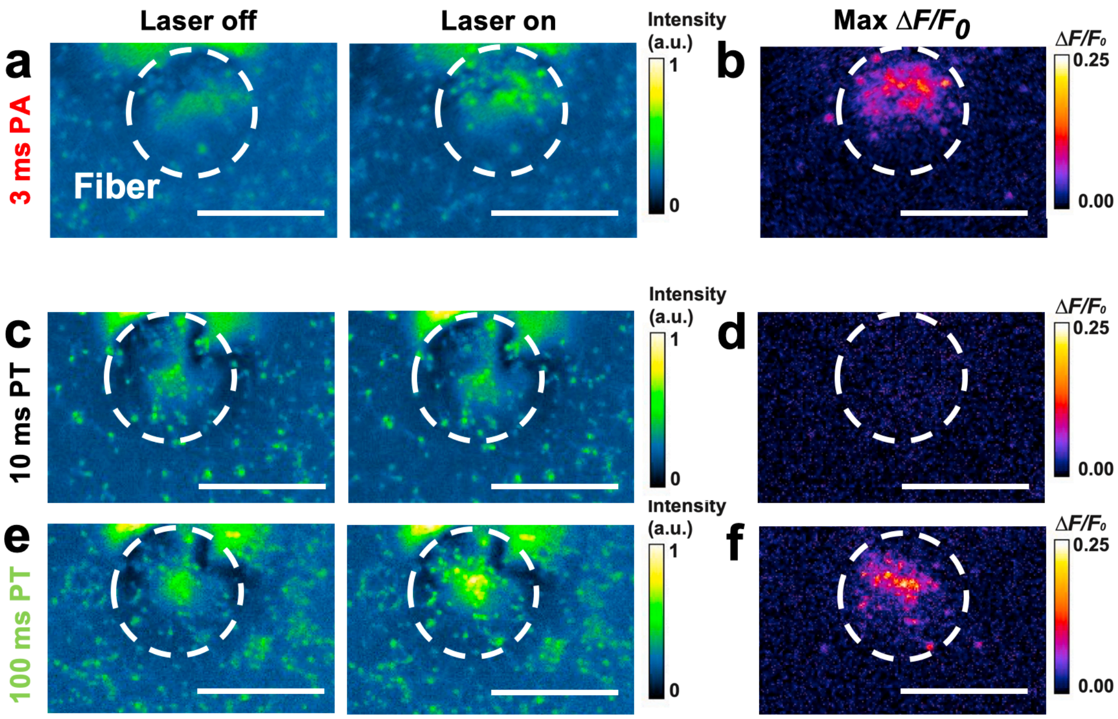


**Figure S2**. **Calcium imaging of PA and PT stimulated neurons.** a-b. representative fluorescence imaging of OGD labeled neuron before / after laser is on. Laser condition: 3 ns pulsed laser, 120 mW, 3 ms burst duration. c-d. representative fluorescence imaging of OGD labeled neuron before / after laser is on. Laser condition: CW laser, 120 mW, 10 ms burst duration. e-f. representative fluorescence imaging of OGD labeled neuron before / after laser is on. Laser condition: CW laser, 120 mW, 100 ms burst duration.

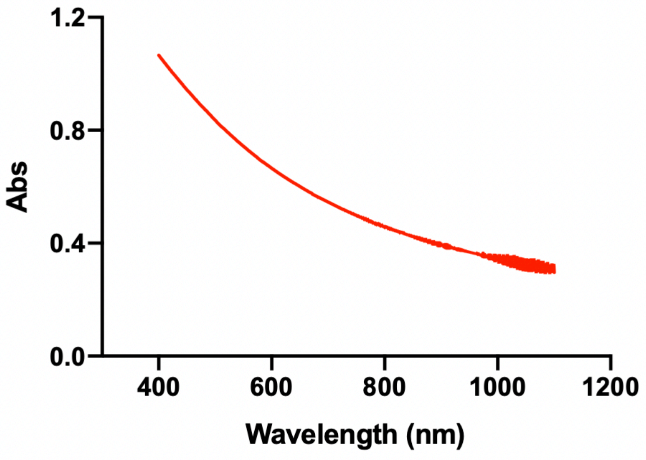


**Figure S3. UV-vis extinction spectrum of candle soot**. The absorption at 1064 nm and 1030 nm are similar.

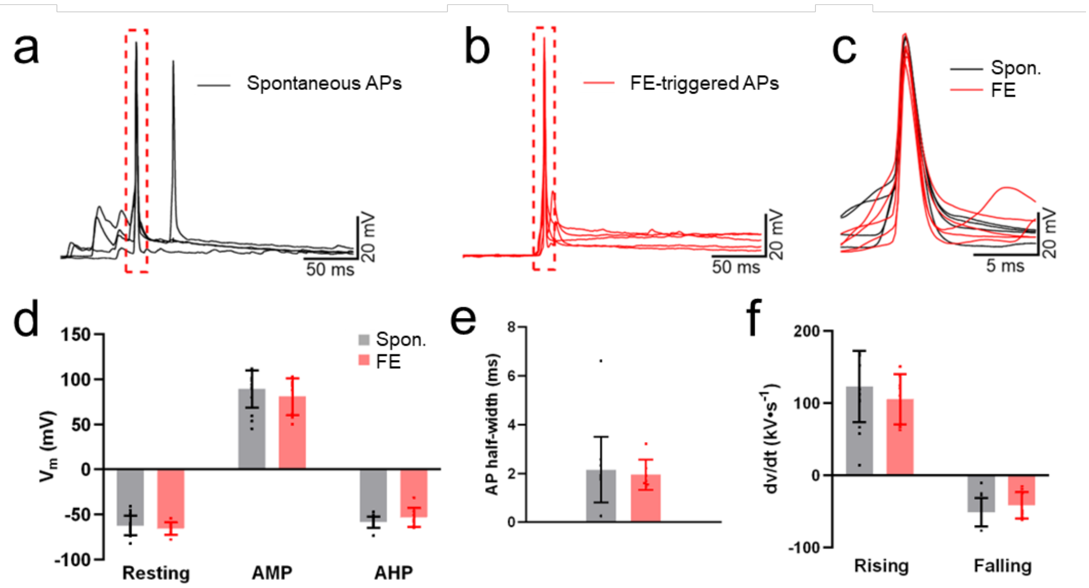


**Figure S4. PA-triggered APs showed similar characteristics as spontaneous APs.** a-b Overlaying individual representative spontaneous (spon.) APs (a) and PA-triggered APs (b) recorded from a neuron in different trials. Red dashed boxes labelled the area further plotted in c. c peaks of the same APs in c and d in an expanded timescale. All traces were normalized to the same baseline and aligned by peaks. d-f Summary of resting membrane potential (Spon. -62.3 ± 10.82 mV, TFOE -65.63 ± 7.157 mV), peak amplitude (AMP, Spon. 89.21 ± 20.55 mV, TFOE 80.78 ± 20.65 mV), afterhyperpolarization (AHP, spon. -58.65 ± 5.959 mV, TFOE -53.46 ± 10.62 mV), AP width at half-maximum peak amplitude (spon. 2.158 ± 1.34 ms, TFOE 1.94 ± 0.62 ms), maximum and minimum rate during rising (spon. 123.0 ± 49.5 kV/s, TFOE 105.4 kV/s) and falling phase (spon. -50.79 ± 19.68 kV/s, TFOE -41.37 ± 18.52 kV/s). Mean ± standard deviation was plotted. N = 15 from 3 cells for Spontaneous and n = 8 from the same 3 cells for TFOE-stimulated trials. No statistically significance was found in *T-tests*.

**Laser pulses vs trigger signals**

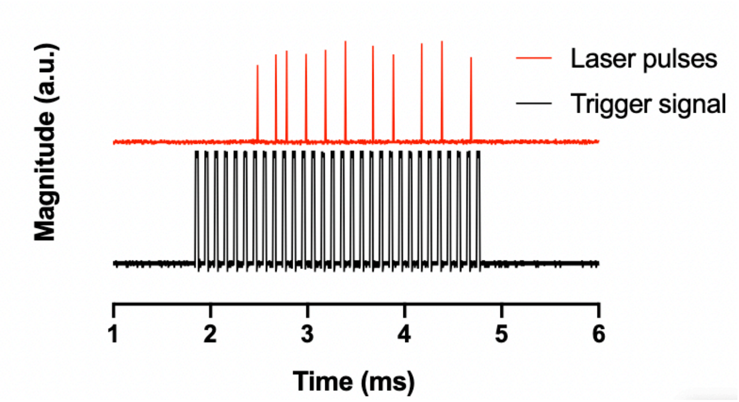


**Figure S5**. Laser pulses versus the trigger signal within 3 ms burst duration. There is a delay of around 0.6 ms after the laser receives the trigger signal.

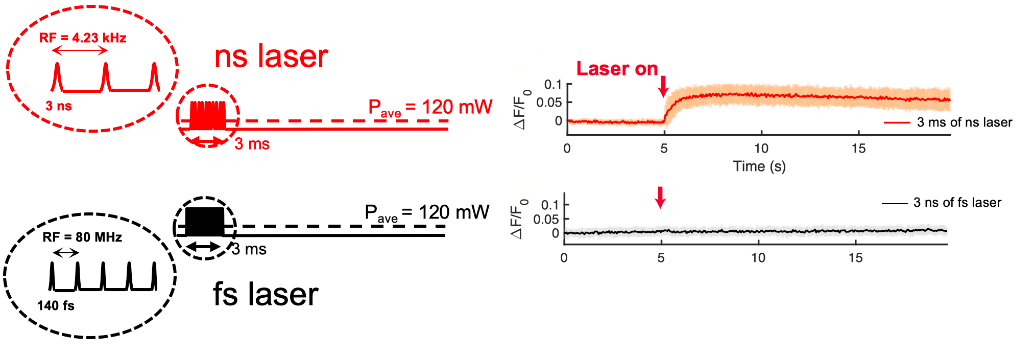


**Figure S6.** Stimulation of primary cortical neurons using a nanosecond pulsed laser (PA condition) and a femtosecond pulsed laser (optocapacitive condition), resplectively. OGD488 fluorescence was recorded. N = 10 for PA condition and N =24 For optocapacitive condition.
